# Stage-specific CHI3L1/YKL-40 signaling controls generation of oligodendrocyte precursor cells through IL13Rα2-mediated ferroptosis

**DOI:** 10.64898/2026.05.08.723841

**Authors:** Xin Yang, Yu-Wen Alvin Huang

**Affiliations:** Department of Molecular Biology, Cell Biology and Biochemistry, Center for Translational Neuroscience, Carney Institute for Brain Science, Brown University, Providence, RI, USA

**Keywords:** Chitinase 3 like protein 1 (CHI3L1/YKL-40), Oligodendrocyte precursor cells (OPCs), Pre-OPC, Ferroptosis, Interleukin-13 receptor subunit alpha-2 (IL13Rα2)

## Abstract

Oligodendrocyte precursor cells (OPCs) are essential for sustaining myelin plasticity and maintaining oligodendroglial homeostasis throughout life. However, the intrinsic signaling mechanisms that regulate OPC generation from neural progenitors remain incompletely understood. Here, we identify chitinase-3-like protein 1 (CHI3L1/YKL-40) as a stage-specific, intrinsic signaling regulator of oligodendroglial lineage progression. Using human iPSC-derived models, we show that CHI3L1 selectively targets a transient IL13Rα2-expressing pre-OPC population, where it induces ferroptosis through suppression of GPX4, thereby limiting OPC generation. In contrast, CHI3L1 does not trigger cell death in established OPCs but instead suppresses their proliferation and differentiation while promoting a reactive, immune-like transcriptional state. Mechanistically, IL13Rα2 mediates CHI3L1-dependent ferroptotic vulnerability at the pre-OPC stage, distinguishing it from parallel signaling pathways that regulate neuronal differentiation. Functionally, this dual-stage regulation constrains the production and maintenance of OPCs, a critical determinant of oligodendroglial homeostasis and myelin plasticity. These findings define an intrinsic signaling axis that couples stage-specific cell fate vulnerability to ferroptotic cell death, thereby controlling OPC pool dynamics with broad implications for cognitive function and disorders involving disrupted myelination.

## Introduction

The generation and maintenance of myelin in the central nervous system (CNS) depend on the continuous production of oligodendrocytes from oligodendrocyte precursor cells (OPCs), a proliferative population distributed throughout both the developing and adult brain (*1–4*). Myelin is dynamically regulated across the lifespan and is essential not only for neural circuit function and learning, but also for adaptive responses to injury (*5, 6*). In the adult CNS, resident OPCs proliferate and differentiate to replenish lost oligodendrocytes, thereby preserving a relatively stable OPC pool. The tight coordination of OPC generation, proliferation, and maturation is therefore fundamental to maintaining myelin integrity and oligodendroglial homeostasis.

While mature OPCs have been extensively studied, much less is known about the transitional stages that precede OPC formation. In particular, the early progenitor state that becomes committed to the oligodendroglial lineage, referred to here as the pre-OPC stage, remains poorly defined (*7*). This stage is likely to represent a critical developmental window during which neural progenitors lose multipotency and acquire oligodendroglial fate (*8*), yet the molecular signals governing this transition are not well understood. Defects at this step could have major downstream consequences by limiting the supply of OPCs available for myelin maintenance and repair (*8*). Because OPC pool size and function are central to myelin plasticity, identifying signaling mechanisms that regulate this early lineage commitment step is essential for understanding oligodendroglial homeostasis and its disruption in disease.

Chitinase-3-like protein 1 (CHI3L1), also known as YKL-40, is a secreted glycoprotein predominantly produced by astrocytes and widely recognized as a biomarker of central nervous system (CNS) inflammation (*9*). Beyond its diagnostic value, accumulating evidence including work from our group demonstrates that CHI3L1 actively regulates cellular responses within the neural environment (*10–13*). We previously showed that astrocyte-derived CHI3L1 suppresses neural stem cell (NSC) proliferation and neuronal differentiation through activation of the CRTH2 receptor and downstream signaling pathways, leading to depletion of the NSC pool and impaired neurogenesis across diverse inflammatory contexts, including neuromyelitis optica (*10*) and Alzheimer’s disease (*11*). Importantly, genetic or pharmacological inhibition of CHI3L1-CRTH2 signaling restores neurogenesis and synaptic functions, and improves functional outcomes, establishing CHI3L1 as a key regulator of neural lineage dynamics (*10–12*). In parallel, CHI3L1 has been shown to engage receptors such as RAGE to promote proinflammatory gliosis (*13*), suggesting that it integrates extracellular stress signals with cell-intrinsic transcriptional and functional responses. Collectively, these studies position CHI3L1 as an active signaling molecule that regulates neural lineage behavior and contributes to pathological remodeling of the CNS. However, whether CHI3L1 directly controls oligodendrocyte lineage progression, particularly during early stages of lineage commitment and myelin-forming cell generation, remains unknown.

In this study, we investigated the role of CHI3L1 in OPC lineage progression using a human iPSC-derived model system (*11, 14*) that enables precise dissection of stage-specific cellular transitions. We identify CHI3L1 as a stage-dependent regulator of oligodendroglial homeostasis. At an early stage of lineage commitment, CHI3L1 selectively targets a transient IL13Rα2-enriched pre-OPC population, inducing ferroptosis through suppression of GPX4 and thereby limiting OPC generation. In contrast, CHI3L1 does not trigger cell death in established OPCs, but instead suppresses their proliferation and differentiation while promoting a reactive, immune-like transcriptional state. This dual mechanism - selective elimination of progenitors coupled with functional reprogramming of committed OPCs - reveals a previously unrecognized regulatory axis that constrains OPC pool dynamics. Together, our findings define a stage-specific signaling pathway that links transient progenitor state vulnerability to ferroptotic cell death and provides a framework for understanding how intrinsic signaling mechanisms regulate oligodendroglial lineage progression and myelin homeostasis.

## RESULTS

### CHI3L1 impairs OPC generation and redirects neural progenitor fate under disease-relevant conditions

We first sought to determine whether CHI3L1 directly impairs OPC generation from neural progenitors, a key process underlying oligodendroglial homeostasis. To place our in vitro studies in a pathophysiologically relevant context, we next defined the extracellular range of CHI3L1 exposure associated with human brain disease. Since CHI3L1 functions as a secreted extracellular signaling molecule, its concentration in cerebrospinal fluid (CSF) provides a clinically relevant estimate of the levels to which neural progenitors and glial lineage cells may be exposed in the diseased CNS. We therefore extracted CSF CHI3L1 measurements from independent clinical studies and performed a meta-analysis across multiple neurological disorders. This analysis corroborated and expanded previous findings by showing that CSF CHI3L1 is elevated across an array of brain diseases, including frontotemporal dementia, multiple sclerosis, and Alzheimer’s disease, with concentrations ranging from below 100 ng/mL in controls to more than 500 ng/mL in severe cases (**Fig. S1A; Table S1**). Guided by this pathophysiologically relevant range, we asked whether elevated CHI3L1 directly alters early neural lineage decisions, particularly the generation of oligodendrocyte precursor cells (OPCs), a critical step in oligodendroglial homeostasis.

To address this, we utilized a human iPSC-derived neural progenitor cell (iNPC) system that recapitulates multipotent neural stem cell (NSC)-like behavior and enables controlled differentiation into neuronal, astroglial, and oligodendroglial lineages (*11, 14*). This platform provides a tractable model to dissect how extracellular signals regulate lineage specification. Consistent with our previous findings, CHI3L1 treatment significantly impaired neurogenesis, as evidenced by reduced DCX-positive cells during differentiation (**Fig. 1A**) and decreased DCX protein levels (**Fig. S1B**). We next examined oligodendrogenesis and found that CHI3L1 markedly reduced the generation of Olig2-positive cells, indicating suppression of oligodendroglial lineage commitment (**Fig. 1B**). This effect was further confirmed by decreased Olig2 protein expression by immunoblotting (**Fig. 1C**). In parallel, CHI3L1 reduced proliferation, as reflected by decreased PCNA levels, and increased astroglial differentiation, indicated by elevated GFAP expression at both the protein (**Fig. S1C**) and immunofluorescence levels (**Fig. S1D**). Together, these results demonstrate that CHI3L1 broadly shifts neural progenitor fate away from neuronal and oligodendroglial lineages and toward astroglial identity, while also suppressing proliferative capacity.

**Figure 1.**
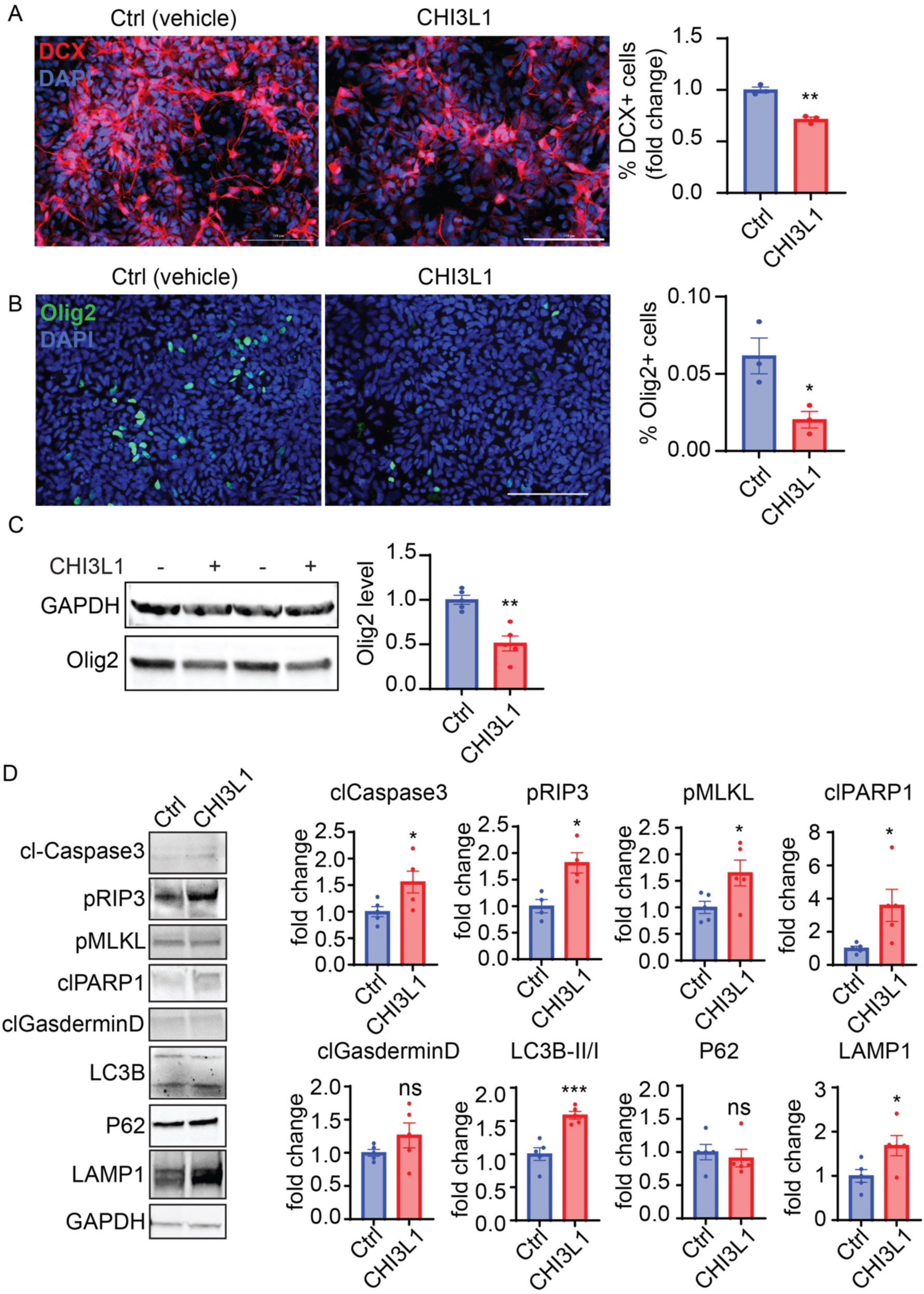
CHI3L1 suppresses neurogenesis and oligodendrogenesis in differentiating human iPSC-derived neural progenitor cells. **(A)** Neurogenesis assay in differentiating human iPSC-derived neural progenitor cells (iNPCs). Representative immunofluorescence images show DCX-positive newly formed neurons (red) in vehicle-and CHI3L1-treated cultures. Nuclei were counterstained with DAPI (blue). Quantification of DCX-positive cells is shown at right as fold change relative to the control condition. CHI3L1 reduced neuronal differentiation. *n* = 3 independent experiments. **(B)** Oligodendrogenesis assay in differentiating iNPCs. Representative immunofluorescence images show Olig2-positive oligodendroglial lineage cells (green) in vehicle- and CHI3L1-treated cultures. Nuclei were counterstained with DAPI (blue). Quantification of the percentage of Olig2-positive cells relative to total DAPI-positive cells is shown at right. CHI3L1 reduced oligodendroglial lineage generation. *n* = 3 independent experiments. **(C)** Immunoblotting analysis of Olig2 protein expression in differentiating iNPCs with or without CHI3L1 treatment. Representative immunoblots are shown at left, and quantification normalized to GAPDH is shown at right. CHI3L1 decreased Olig2 protein levels. *n* = 4 blots from 3 independent experiments. **(D)** Immunoblotting analysis of cell death- and stress-related pathway markers in differentiating iNPCs following CHI3L1 treatment. Representative immunoblots for cleaved caspase-3, phosphorylated RIP3, phosphorylated MLKL, cleaved PARP1, cleaved gasdermin D, LC3B, p62, and LAMP1 are shown at left, with quantification normalized to GAPDH shown at right. CHI3L1 increased markers associated with apoptosis, necroptosis, and autophagy-related stress responses in differentiating iNPCs. *n* = 5 blots from 3 independent experiments. Data are presented as mean ± SEM. Statistical significance was determined by unpaired *t* test, Ctrl vs. CHI3L1. * p < 0.05, ** p < 0.01, *** p < 0.001; ns, not significant.

The decrease in OPC generation after CHI3L1 treatment could result from either defective lineage specification or increased cell death during differentiation. Because cell death is a well-recognized feature of oligodendroglial development and can influence lineage output (*16*), we next asked whether CHI3L1 activates death-related pathways in differentiating neural progenitors. Immunoblotting showed increased levels of cleaved caspase-3 and cleaved PARP, consistent with apoptotic signaling, as well as increased phosphorylated RIP3 and MLKL, indicative of necroptosis-associated signaling, and elevated LC3B-II, consistent with autophagy-associated stress (**Fig. 1D**). In contrast, cleaved gasdermin D was not significantly changed. Together, these findings indicate that CHI3L1 triggers a broad cellular stress response during early differentiation, raising the possibility that activation of specific cell death programs contributes to the loss of OPC generation.

To distinguish among senescence, cell death, and altered lineage progression as potential mechanisms underlying the CHI3L1 phenotype, we next examined transcriptional changes in differentiating neural progenitors. To further define the underlying mechanisms, we examined transcriptional changes associated with cell cycle regulation, senescence, and lineage specification. Expression of canonical senescence markers (p16, p21, and p27) (*17, 18*) was not significantly altered (**Fig. S1E**), arguing against a primary role for senescence. In contrast, expression of cell death-related genes, including TP53 and CASP9, was increased, consistent with activation of apoptotic pathways. Analysis of lineage-associated genes revealed that early oligodendroglial markers such as PDGFRα and Olig1 were largely unchanged, whereas NKX2.2, a key regulator of oligodendrocyte specification, was significantly reduced. Neuronal differentiation was similarly impaired, as indicated by decreased TUBB3 expression. Notably, SOX2 expression was increased, suggesting that CHI3L1 maintains cells in a progenitor-like state and prevents progression toward differentiated neuronal and oligodendroglial fates.

Collectively, these findings demonstrate that CHI3L1 suppresses OPC generation by simultaneously impairing lineage specification and inducing cellular stress responses during early differentiation. These observations raise the key mechanistic question of whether CHI3L1 targets a specific stage within the oligodendroglial lineage that is particularly vulnerable to its effects.

### Blocking IL13Rα2 and ferroptosis ameliorates OPC generation

Given that CHI3L1 suppresses OPC generation, we next sought to identify the receptor(s) mediating this effect. Multiple receptors have been proposed for CHI3L1 signaling, including CD44, CRTH2, GAL3, IL13Rα2, RAGE, and TMEM219 (*9, 19*). While our previous work identified CRTH2 as a key mediator of CHI3L1-induced neurogenesis deficits and RAGE as a regulator of astrocyte-intrinsic responses (*10, 11*), it remains unclear which receptor(s) govern CHI3L1 effects on oligodendroglial lineage progression.

To address this, we first performed a systematic reanalysis of the published single-cell RNA-seq dataset GSE75330 to examine receptor expression across oligodendroglial lineage states. All known CHI3L1 receptors were detectable across the lineage, spanning OPCs to mature oligodendrocytes (**Fig. 2A**). We then validated these findings in our human iPSC-derived system by qPCR, confirming expression of CD44, CRTH2, GAL3, IL13RA2, RAGE, and TMEM219 in iNPCs, iOPCs, and induced oligodendrocytes (iOLs) (**Fig. S2A**). While most receptors were expressed at comparable levels across stages, GAL3 and TMEM219 showed increased expression during oligodendroglial differentiation. Cell identity across stages was confirmed by expected marker dynamics, including downregulation of PAX6, sustained Olig2 expression, and induction of MBP in iOLs (**Fig. S2B**). Independent analysis of mouse brain RNA-seq data further supported these patterns and revealed enrichment of TMEM219 in oligodendrocyte-lineage populations (**Fig. S2C**).

**Figure 2.**
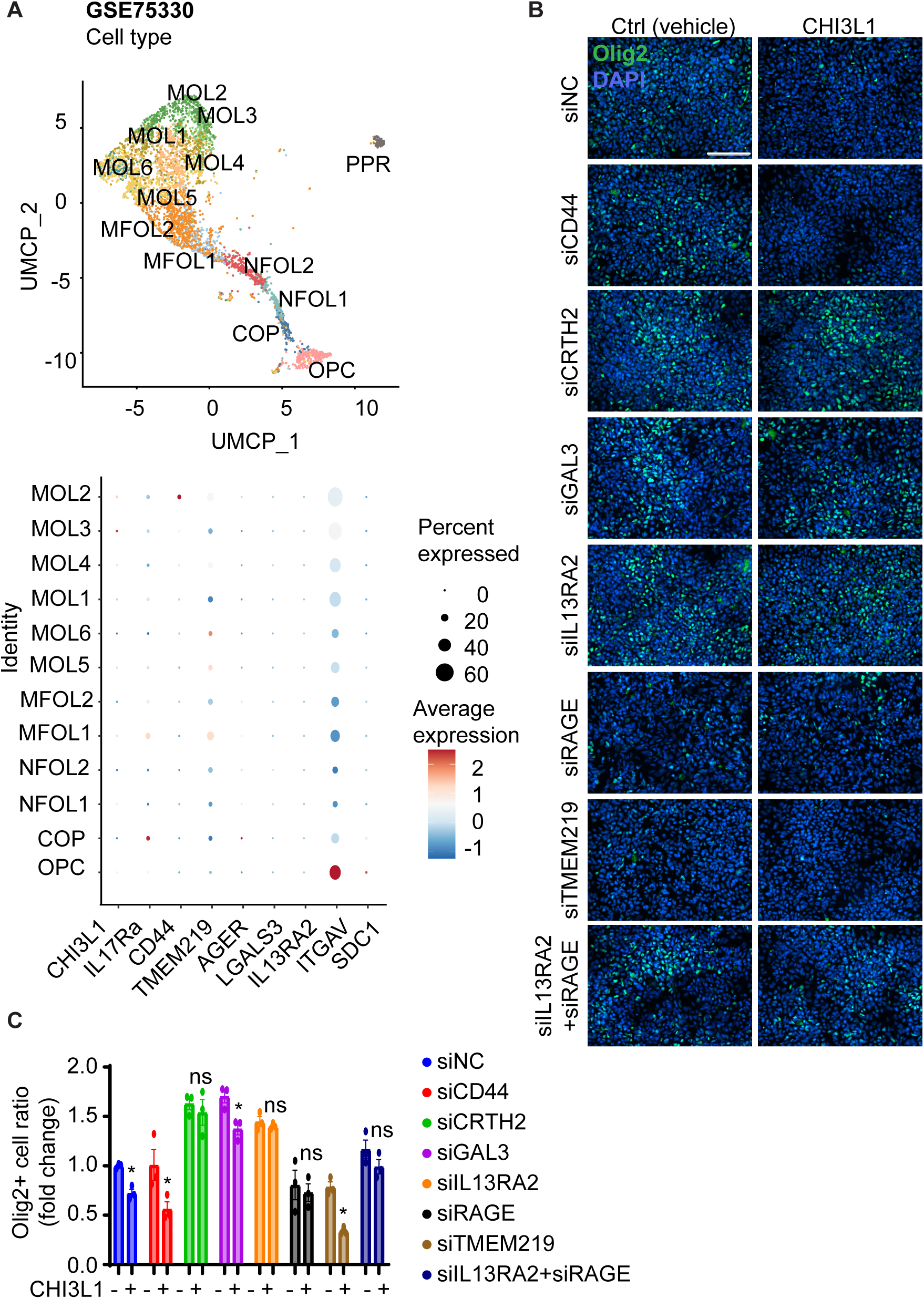
IL13Rα2 mediates CHI3L1-dependent suppression of oligodendroglial lineage commitment. **(A)** Reanalysis of the published single-cell RNA-seq dataset GSE75330 to examine the expression of known CHI3L1 receptors across oligodendroglial lineage stages. UMAP visualization shows transcriptionally defined populations, including OPCs, committed oligodendrocyte precursors (COPs), newly formed oligodendrocytes (NFOL1/2), myelin-forming oligodendrocytes (MFOL1/2), and mature oligodendrocytes (MOL1–6), as well as perivascular cells (PPR). Dot plot analysis depicts the relative expression (color scale) and percentage of cells expressing (dot size) CHI3L1 and its reported receptors (IL17RA, CD44, TMEM219, AGER, LGALS3, IL13RA2, ITGAV, SDC1) across these populations. **(B)** Immunofluorescence staining of Olig2 (green) in differentiating human iPSC-derived neural progenitor cells (iNPCs) treated with control (vehicle) or CHI3L1 following siRNA-mediated knockdown of the indicated CHI3L1 receptors (siNC, siCD44, siCRTH2, siGAL3, siIL13RA2, siRAGE, siTMEM219, and combined siIL13RA2+siRAGE). Nuclei were counterstained with DAPI (blue). Representative images show the effects of receptor-specific knockdown on CHI3L1-induced reduction of oligodendroglial lineage cells. **(C)** Quantification of Olig2-positive cells from (B), shown as fold change relative to control conditions. Knockdown of IL13RA2 selectively rescues the reduction in Olig2-positive cells induced by CHI3L1, whereas knockdown of other receptors shows minimal or partial effects. Combined knockdown of IL13RA2 and RAGE does not further enhance rescue compared to IL13RA2 alone. *n* = 3 independent experiments. Data are presented as mean ± SEM. Statistical significance was determined by one-way ANOVA with post-hoc analyses and selected comparisons shown (without or with CHI3L1); * p < 0.05, ns, not significant.

We next tested the functional contribution of each receptor using siRNA-mediated knockdown in differentiating iNPCs. Efficient knockdown of each receptor was confirmed by qPCR (**Fig. S2D**). Under basal conditions, knockdown of CD44, CRTH2, or IL13Rα2 modestly increased the generation of Olig2-positive cells, suggesting that these receptors may exert tonic inhibitory effects on oligodendroglial lineage output (**Fig. 2B-C**). Upon CHI3L1 treatment, however, a more selective pattern emerged: knockdown of IL13Rα2 robustly rescued the CHI3L1-induced reduction in Olig2-positive cells, whereas knockdown of CRTH2 provided partial rescue, and depletion of other receptors had minimal effect (**Fig. 2B-C**). These results identify IL13Rα2 as the primary receptor mediating CHI3L1-dependent suppression of OPC generation.

To determine whether receptor specificity extends to lineage context, we examined neurogenesis under the same conditions. Consistent with our previous findings (*11*), knockdown of CRTH2, but not IL13Rα2, rescued CHI3L1-induced suppression of DCX-positive neurons (**Fig. S2E**), indicating that CHI3L1 engages distinct receptor pathways to regulate neuronal versus oligodendroglial lineage outcomes.

Together, these findings establish receptor-specific control of CHI3L1 signaling, with IL13Rα2 selectively mediating its inhibitory effects on OPC generation. Given our earlier observation that CHI3L1 induces broad stress and cell death-associated signaling, these results further suggest that IL13Rα2 may couple CHI3L1 signaling to a specific downstream cell death mechanism that limits OPC production.

### CHI3L1 induces IL13Rα2-dependent ferroptosis to suppress OPC generation

Ferroptosis has emerged as a key regulator of oligodendrocyte lineage viability, with accumulating evidence demonstrating its role in OPC loss and impaired myelination under pathological conditions (*20–22*). Given our observation that CHI3L1 activates broad stress and cell death–associated pathways, we next asked whether a specific cell death modality mediates its inhibitory effect on OPC generation.

To address this, differentiating iNPCs were pre-treated with inhibitors targeting major cell death pathways, including apoptosis (Z-VAD-FMK), necroptosis (necrostatin-1), ferroptosis (ferrostatin-1), and pyroptosis (disulfiram), followed by CHI3L1 exposure. Strikingly, only ferroptosis inhibition significantly rescued the reduction in Olig2-positive cells induced by CHI3L1 (**Fig. 3A**), whereas inhibition of other pathways had minimal effect. Consistent with this specificity, ferroptosis inhibition did not substantially restore neurogenesis (**Fig. S2F**), indicating that CHI3L1 regulates oligodendroglial and neuronal lineage outcomes through distinct mechanisms. These findings identify ferroptosis as the dominant cell death pathway underlying CHI3L1-mediated suppression of OPC generation.

**Figure 3.**
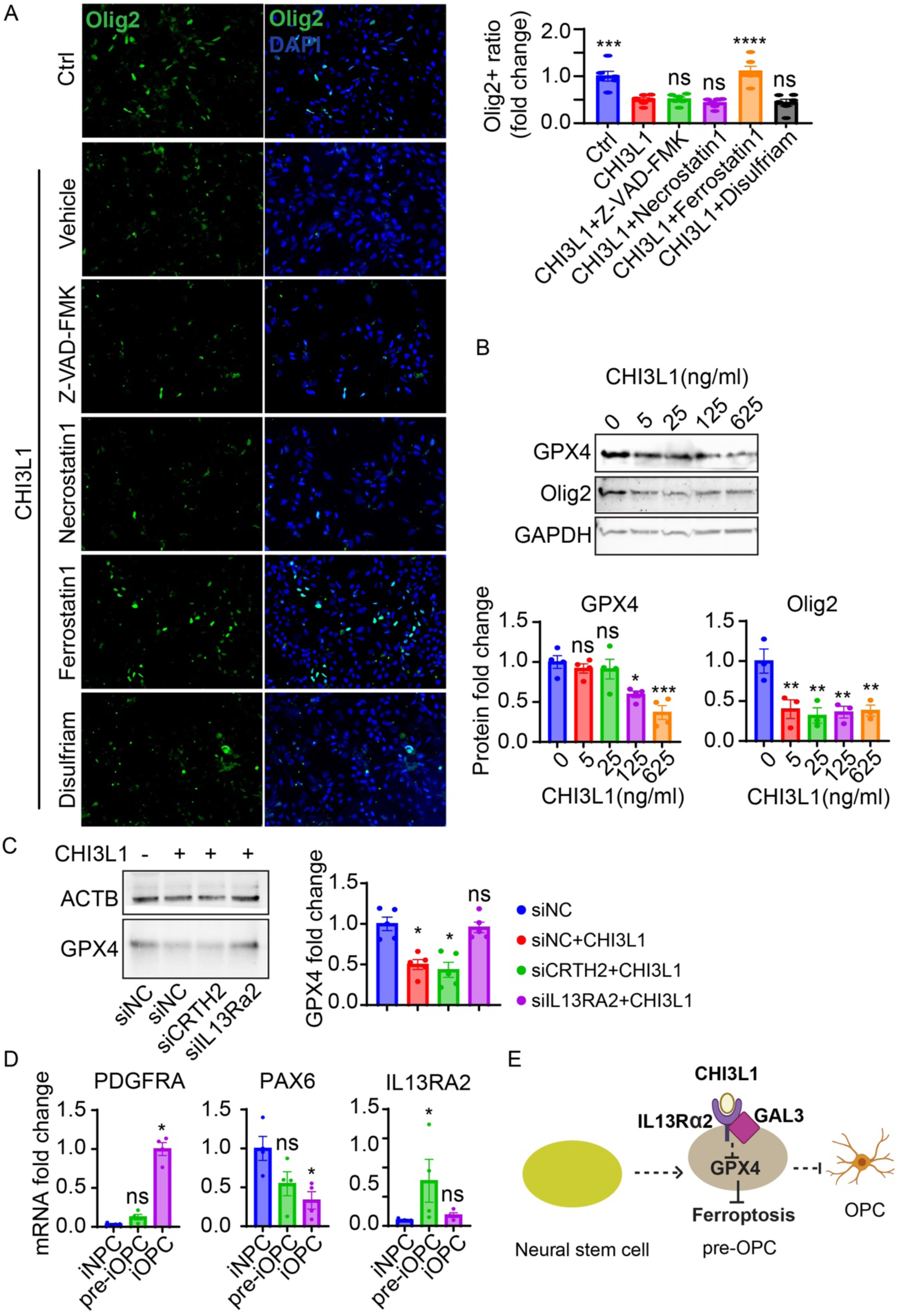
CHI3L1 induces IL13Rα2-dependent ferroptosis to suppress OPC generation. **(A)** Immunofluorescence staining of Olig2 (green) in differentiating human iPSC-derived neural progenitor cells (iNPCs) pre-treated with the indicated cell death inhibitors, including Z-VAD-FMK (apoptosis inhibitor), necrostatin-1 (necroptosis inhibitor), ferrostatin-1 (ferroptosis inhibitor), and disulfiram (pyroptosis inhibitor), in the presence or absence of CHI3L1. Nuclei were counterstained with DAPI (blue). Representative images are shown, with quantification of Olig2-positive cells at right. Ferrostatin-1 selectively rescues CHI3L1-induced reduction in oligodendroglial lineage cells. *n* = 3 independent experiments. **(B)** Immunoblotting analysis of GPX4 and Olig2 expression in differentiating iNPCs treated with increasing concentrations of CHI3L1 (0–625 ng/mL). Representative immunoblots are shown above, with quantification normalized to GAPDH shown below. CHI3L1 reduces GPX4 and Olig2 protein levels in a dose-dependent manner. *n* = 4 independent experiments. **(C)** Immunoblotting analysis of GPX4 expression in iNPCs following siRNA-mediated knockdown of the indicated receptors (siNC, siCRTH2, siIL13RA2) in the presence or absence of CHI3L1. Representative immunoblots are shown at left, with quantification normalized to β-actin (ACTB) shown at right. Knockdown of IL13RA2, but not CRTH2, rescues CHI3L1-induced reduction of GPX4. *n* = 5 independent experiments. **(D)** qPCR analysis of lineage progression and receptor expression in iNPCs, early differentiating iNPCs (pre-OPCs), and iOPCs. Expression of PDGFRA, PAX6, and IL13RA2 is shown as fold change, revealing transient enrichment of IL13RA2 during early lineage commitment. *n* = 4 independent experiments. **(E)** Schematic model illustrating CHI3L1 signaling through IL13Rα2 to suppress GPX4 and induce ferroptosis in a transient pre-OPC population, thereby limiting OPC generation from neural progenitors. Data are presented as mean ± SEM. Statistical significance was determined by one-way ANOVA with post-hoc analyses; significance indicated as compared to the control conditions (blue bars) in (B-D), or to the CHI3L1 condition (red bar) in (A); * p < 0.05, ** p < 0.01, *** p < 0.001, **** p < 0.0001; ns, not significant.

We next investigated whether CHI3L1 directly engages the ferroptotic machinery. Immunoblotting revealed that CHI3L1 reduced expression of glutathione peroxidase 4 (GPX4), a central inhibitor of ferroptosis (*23*), in a dose-dependent manner, accompanied by decreased Olig2 levels (**Fig. 3B**). Importantly, the CHI3L1-induced reduction in GPX4 was selectively reversed by ferroptosis inhibition, but not by inhibitors of other cell death pathways (**Fig. S3A**), further supporting a ferroptosis-specific mechanism.

Because ferroptosis is driven by iron-dependent oxidation of polyunsaturated lipids in cellular membranes (*24, 25*), we next asked whether antioxidant treatment could counteract the effects of CHI3L1. To test this, differentiating iNPCs were pre-incubated with N-acetylcysteine (NAC), a general antioxidant known to protect OPCs from oxidative injury (*26*), before CHI3L1 exposure. NAC pre-treatment restored GPX4 protein levels and prevented the CHI3L1-induced reduction in Olig2 expression (**Fig. S3B-C**). These findings support the interpretation that CHI3L1 suppresses OPC generation through oxidative stress–dependent ferroptotic signaling.

We then examined whether this pathway is mediated through IL13Rα2, the receptor identified above as a key regulator of CHI3L1-dependent OPC suppression. Knockdown of IL13Rα2 abolished CHI3L1-induced downregulation of GPX4, whereas CRTH2 knockdown had no such effect (**Fig. 3C**). These results demonstrate that IL13Rα2 is specifically required to couple CHI3L1 signaling to ferroptotic machinery.

Finally, we asked whether IL13Rα2 expression is dynamically regulated during lineage progression, potentially defining a vulnerable cell state. Although IL13Rα2 levels were comparable between bulk iNPC and iOPC populations (**Fig. S2A**), qPCR analysis across differentiation stages revealed a transient upregulation of IL13Rα2 in early differentiating progenitors (**Fig. 3D**; pre-iOPCs, 3 days post-differentiation). This expression peak coincided with the transition from PAX6-positive neural progenitors to PDGFRA-positive oligodendroglial progenitors, suggesting the existence of a pre-OPC state with heightened sensitivity to CHI3L1 signaling.

Together, these findings support a model in which CHI3L1, acting through IL13Rα2, suppresses GPX4 and induces ferroptosis in a transiently vulnerable pre-OPC population, thereby limiting OPC generation (**Fig. 3E**).

### CHI3L1 switches OPC from proliferation and differentiation to inflammation

CHI3L1 suppresses OPC proliferation and induces inflammatory reprogramming without activating cell death pathways (*12, 13, 27*), but the cellular mechanisms underlying this effect - whether through cell death, senescence, or altered proliferative capacity - remain unclear. To address this, we examined the direct impact of CHI3L1 on committed oligodendroglial populations using purified induced oligodendrocyte precursor cells (iOPCs) and newly formed induced oligodendrocytes (iOLs). iOPCs were generated as described and differentiated into iOLs by PDGF-AA withdrawal with supplementation of T3 and clemastine (*14*). Cell identity was validated by marker expression: iOPCs exhibited high levels of Olig2 (**Fig. S2B**) and PDGFRα (**Fig. 4A**), whereas differentiated iOLs retained Olig2 but showed loss of PDGFRα and acquisition of mature oligodendrocyte markers including MBP and O4 (**Fig. 4A, Fig. S2B**).

**Figure 4.**
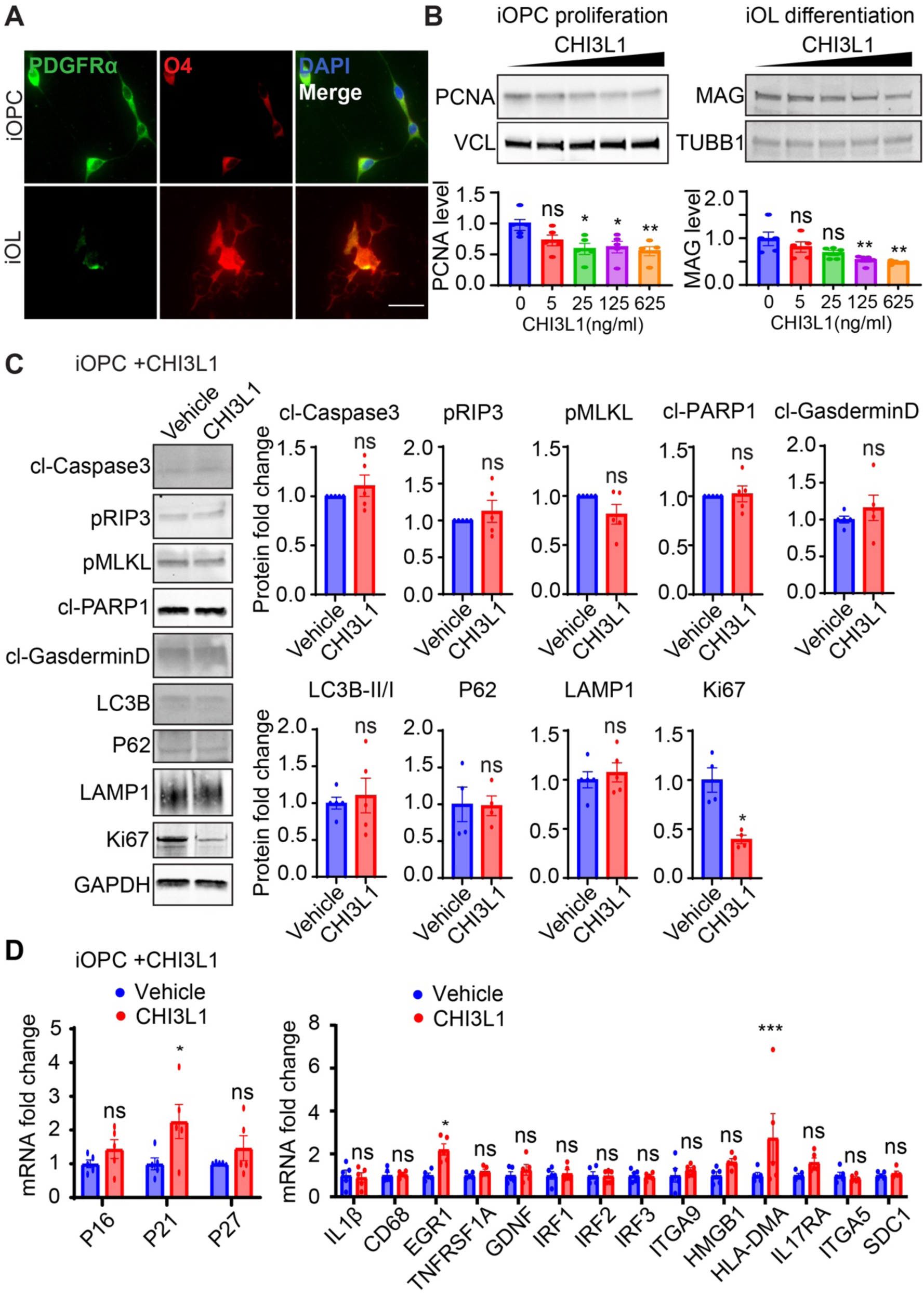
CHI3L1 impairs proliferation and induces inflammatory reprogramming in OPCs without activating cell death pathways. **(A)** Immunofluorescence staining of PDGFRα (green) and O4 (red) in induced oligodendrocyte precursor cells (iOPCs) and newly formed induced oligodendrocytes (iOLs). Nuclei were counterstained with DAPI (blue), confirming stage-specific identity of OPCs and differentiating oligodendrocytes. **(B)** Top: Experimental schematic illustrating CHI3L1 treatment during iOPC proliferation (left) and differentiation into iOLs (right). Bottom left: Immunoblotting analysis of proliferation marker PCNA and differentiation marker MAG in iOPCs treated with increasing concentrations of CHI3L1. Bottom right: Quantification of PCNA and MAG protein levels normalized to vinculin (VCL) and β-tubulin (TUBB1), respectively. CHI3L1 reduces OPC proliferation and impairs differentiation in a dose-dependent manner. *n* = 4 independent experiments. **(C)** Immunoblotting analysis of cell death–associated markers in iOPCs treated with or without CHI3L1 (500 ng/mL). Representative blots and quantification show no significant activation of apoptosis (cleaved caspase-3, cleaved PARP1), necroptosis (pRIP3, pMLKL), pyroptosis (cleaved gasdermin D), or autophagy-associated markers (LC3B, p62, LAMP1). In contrast, the proliferation marker Ki67 is reduced following CHI3L1 treatment. Protein levels were normalized to GAPDH. *n* = 4 independent experiments. **(D)** qPCR analysis of senescence-associated genes (P16, P21, P27) and inflammatory or stress-response genes (IL1β, CD68, EGR1, TNFRSF1A, GDNF, IRF1–3, ITGA9, HMGB1, HLA-DMA, IL17RA, ITGA5, SDC1) in iOPCs following CHI3L1 treatment. CHI3L1 selectively induces inflammatory gene expression without robust activation of senescence-associated pathways. *n* = 4 independent experiments. Data are presented as mean ± SEM. Statistical significance was determined by unpaired *t* test in (C) one-way ANOVA with post-hoc analyses and the selected comparisons shown (Vehicle vs. CHI3L1); * p < 0.05, ** p < 0.01, *** p < 0.001; ns, not significant.

We next assessed how CHI3L1 influences OPC proliferation and differentiation. Treatment with increasing concentrations of CHI3L1 resulted in a dose-dependent reduction in proliferation, as indicated by decreased PCNA levels, and impaired differentiation, reflected by reduced expression of myelin-associated glycoprotein (MAG), with effects observed at concentrations of 25–125 ng/mL (**Fig. 4B**). Based on these findings, we used a disease-relevant concentration (500 ng/mL) for subsequent mechanistic studies. Notably, in contrast to our findings in iNPCs, CHI3L1 treatment of iOPCs did not activate canonical cell death pathways. Immunoblotting revealed no significant changes in markers of apoptosis (cleaved caspase-3, cleaved PARP), necroptosis (pRIP3, pMLKL), pyroptosis (cleaved gasdermin D), autophagy (LC3B, p62, LAMP1), or ferroptosis (GPX4) (**Fig. 4C**). Instead, CHI3L1 significantly reduced Ki67 expression (**Fig. 4C**), consistent with decreased PCNA levels and indicating a primary effect on proliferation rather than survival. These findings reveal a stage-dependent shift in CHI3L1 function, from inducing ferroptotic cell death in progenitors to suppressing proliferation and differentiation in committed OPCs.

To determine whether CHI3L1 induces a senescence-like state in iOPCs, we next examined expression of canonical cell cycle inhibitors and senescence markers. While p21 expression was modestly increased, p16 and p27 remained unchanged following CHI3L1 treatment (**Fig. 4D**). Analysis of additional senescence-associated genes, including IL1β and HMGB1 (*17, 18*), showed minimal changes, with only a trend toward increased HMGB1 expression. These results argue against robust induction of a classical senescence program in OPCs.

Given the absence of strong senescence or cell death responses, we next asked whether CHI3L1 promotes an alternative transcriptional state. qPCR analysis revealed that CHI3L1 significantly upregulated early growth response 1 (EGR1) and major histocompatibility complex class II DM alpha (HLA-DMA) (**Fig. 4D**). These changes are consistent with a shift toward a reactive or immunomodulatory phenotype. EGR1 has been implicated in limiting OPC differentiation following white matter injury (*28*), while HLA-DMA is part of the MHC class II antigen presentation pathway (*29*) and is associated with immune activation in oligodendroglial cells (*30*). In demyelinating conditions such as experimental autoimmune encephalomyelitis (EAE) and multiple sclerosis (MS), OPCs can upregulate MHC molecules and interact with immune cells, contributing to a pro-inflammatory microenvironment that impairs remyelination (*31–33*). Thus, CHI3L1-induced upregulation of HLA-DMA suggests a potential mechanism by which elevated CHI3L1 may promote immune-associated dysfunction in oligodendroglial cells and contribute to remyelination failure *in vivo* (*28, 34, 35*).

## Discussion

Our study identifies CHI3L1/YKL-40 as a stage-dependent regulator of oligodendroglial lineage homeostasis. Using human iPSC-derived neural progenitor and oligodendroglial models, we show that CHI3L1 suppresses OPC generation from neural progenitors, reduces OPC proliferation, impairs oligodendrocyte differentiation, and promotes a reactive inflammatory state in committed OPCs. Mechanistically, CHI3L1 acts through distinct pathways depending on cellular stage: in differentiating progenitors, it targets a vulnerable pre-OPC state through IL13Rα2-dependent ferroptosis, whereas in established OPCs it suppresses proliferation and differentiation without inducing overt cell death. These findings extend CHI3L1 beyond its established role as a biomarker of CNS inflammation and position it as an active extracellular signal capable of reshaping oligodendroglial lineage output. Given the importance of OPC generation and myelin plasticity for cognitive function, this pathway may have broad relevance to neurocognitive disorders across neurodevelopmental and neurodegenerative contexts.

OPC differentiation is essential for maintaining the oligodendrocyte pool and supporting adaptive myelination throughout life. Under homeostatic conditions, resident OPCs replenish lost oligodendrocytes through coordinated proliferation and differentiation (4), whereas failure of OPC recruitment, survival, or maturation contributes to impaired myelin repair in demyelinating and neurodegenerative diseases (*36–38*). Cell death is increasingly recognized as a key mechanism shaping oligodendroglial lineage output, particularly during developmental transitions and in pathological environments. In our system, CHI3L1 reduced Olig2-positive OPC generation while inducing stress and death-associated pathways in differentiating neural progenitors. In contrast, CHI3L1 did not activate cell death markers in committed OPCs, but instead reduced PCNA and Ki67 and suppressed MAG expression. These data support a model in which CHI3L1 limits oligodendroglial homeostasis through two separable mechanisms: depletion of newly specified OPCs during early lineage progression and functional suppression of committed OPCs during proliferation and differentiation.

A major conceptual advance of this work is the identification of ferroptosis as a stage-specific mechanism by which CHI3L1 restricts OPC generation. Ferroptosis is particularly relevant to the oligodendrocyte lineage because oligodendroglial cells accumulate iron during differentiation and depend heavily on lipid metabolism for myelin production, creating vulnerability when antioxidant defenses are compromised (*20–22*). We found that CHI3L1 suppresses GPX4, a central ferroptosis-protective factor, and that ferrostatin-1 or NAC restores GPX4 and Olig2 expression. Importantly, CHI3L1-induced GPX4 suppression required IL13Rα2 but not CRTH2, distinguishing this pathway from the CRTH2-dependent neurogenesis defect we previously described (*10, 11*). The transient, stage-specific upregulation of IL13Rα2 during early oligodendroglial commitment suggests that pre-OPCs represent a discrete vulnerability window in which extracellular inflammatory cues can determine whether neural progenitors successfully enter the OPC lineage. This provides a potentially pioneering mechanism linking disease-associated CHI3L1 elevation to impaired OPC homeostasis and subsequent deficits in myelinating differentiation.

Several caveats should be considered. First, our study primarily uses human iPSC-derived 2D culture systems, which allow mechanistic dissection but do not fully capture the cellular complexity, regional architecture, or inflammatory milieu of the CNS in vivo. Second, although we used disease-relevant CHI3L1 concentrations informed by clinical CSF studies, local tissue concentrations and cell-specific exposure in diseased brain regions remain difficult to define. Third, IL13Rα2 marks a transient pre-OPC-like state in our system, but additional single-cell, spatial, or lineage-tracing studies will be needed to determine whether IL13Rα2 defines an equivalent transitional OPC-generating population in vivo. Finally, future studies using co-culture, organoid, demyelination, or conditional genetic models will be important to validate how CHI3L1 signaling affects oligodendroglial development and repair in more physiological contexts.

Therapeutically, our findings suggest that targeting CHI3L1 signaling may preserve oligodendroglial homeostasis at multiple levels. Inhibiting CHI3L1, blocking IL13Rα2-dependent ferroptosis, or restoring antioxidant protection may help protect pre-OPCs and sustain OPC generation, while reducing CHI3L1-driven inflammatory reprogramming may improve the proliferative and differentiating capacity of committed OPCs. These strategies may also enhance approaches aimed at endogenous NSC mobilization or transplantation of iPSC-derived NSCs or OPCs, where a CHI3L1-rich pathological environment could otherwise limit survival, expansion, and myelinating potential. Together, our data reveal CHI3L1 as an active regulator of oligodendroglial lineage vulnerability and identify the CHI3L1–IL13Rα2–GPX4 axis as a candidate therapeutic target for supporting myelin repair and cognitive resilience.

## MATERIALS AND METHODS

### Human iPSC culturing and trans-differentiation into neural precursor cells (iNPCs) for in vitro Oligodendrogenesis

Human iPSCs, the control reference KOLF2.1J line sourced from the NIH-funded iPSC Neurodegenerative Disease Initiative (*15*) were provided by the Jackson Laboratory. KOLF2.1J iPSCs were cultured in mTeSR Plus medium (STEMCELL Technologies, #05825) and expanded in Stem Flex medium (Fisher Scientific, #A3349401) in the stem cell-qualified Membrane Matrix (Fisher Scientific, #A1413201) coated 6-well plates. Neural precursor cells (iNPCs) were derived using a chemically defined dual-SMAD inhibitor medium (STEMCELL Technologies, #08582) over 7 days, as we described previously (*11, 14*). Subsequently, iNPCs were maintained in proliferation (SMAD inhibitor-containing) medium. The medium was refreshed every two days.

To investigate whether CHI3L1 contributes to neurogenesis and Oligodendrogenesis, 2 × 10^5 iNPCs were seeded onto Matrigel-coated 12-well plate (western blotting and q-PCR) or coverslips in a 24-well plate (immunostaining). Medium was switched to comprising Neurobasal medium (Gibco, #21103049) with 2% B27 supplement (Gibco, #17504044), 1% N2 supplement (Gibco, #17502048), FGF-2 (20 ng/ml; PeproTech, #K1606), EGF (20 ng/ml; PeproTech,#A2306), and 2 mM l-glutamine (Gibco, #25030-081), 1uM SAG(MCE, HY-12848), and 10ng/ml PDGF-AA (MCE, HY-P70598) at 37°C in a 5% CO_2_ incubator for indicated time with or without CHI3L1 and chemicals treatment. Cells were harvested for q-PCR, western blotting, or fixed with 4% PFA for immunostaining.

iOPCs were chemically induced as previously described (*14*). Briefly, iNPCs were seeded into Matrigel-coated 6-well plates under iOPC induction medium: D/F-12 containing 1% B27 supplement (Gibco, #17504044), 1% N2 supplement (Gibco, #17502048), FGF-2 (20 ng/ml; PeproTech, #K1606), 1 μM SAG (MCE, HY-12848), 5 μg/ml N-acetylcysteine (NAC; Sigma-Aldrich, #A8199), 5 μM forskolin (MCE, #HY-15371), and 10ng/ml PDGF-AA (MCE, HY-P70598). OPCs) were expanded in proliferation medium containing PDGF-AA. To initiate differentiation, the medium was switched to differentiation medium for 7 days. This medium was identical to the proliferation medium but lacked PDGF-AA and was further supplemented with 1 µM cAMP, 100 nM clemastine, 100 nM rolipram, and 100 ng/mL 3,3’,5-triiodo-L-thyronine (T3). The medium was replaced every 48 hours.

### Immunofluorescence staining of iNPC differentiation in vitro

For immunocytochemistry, cultured cells were fixed with 4% paraformaldehyde (PFA) in PBS for 20 minutes at room temperature. Following fixation, cells were washed with PBS and then permeabilized and blocked for 1 hour at room temperature in a blocking solution consisting of 3% donkey serum and 0.1% Triton X-100 in PBS.

Cells were incubated overnight at 4°C with primary antibodies diluted in the blocking solution. The next day, cells were washed three times with PBS and incubated with appropriate fluorophore-conjugated secondary antibodies for 2 hours at room temperature. Nuclei were counterstained with 4’,6-diamidino-2-phenylindole (DAPI). Stained cells were visualized and imaged using either a laser scanning confocal microscope or a BioTek automated microscope, with all acquisition settings maintained consistently across samples.

The following primary antibodies and dilutions were used: mouse anti-GFAP (1:1000; Antibodies Incorporated, #75-240), chicken anti-Tuj1 (1:1000; GeneTex, #GTX85469), chicken anti-Olig2, rabbit anti-CHI3L1 (1:1000; Abcam, #ab77528 and Thermo Fisher Scientific, #PA5-43746), rat anti-PDGFRα, mouse anti-O1, mouse anti-DCX (1:400; Santa Cruz Biotechnology, #sc-271390), rabbit anti-PAX6 (1:1000; STEMCELL Technologies, #60094), and guinea pig anti-Map2 (1:1000; Synaptic Systems, #188 004). The following secondary antibodies (all raised in donkey) were used at a 1:500 dilution (or as optimized): anti-mouse Alexa Fluor 488, 568, and 647; anti-rabbit Alexa Fluor 488, 568, and 647; anti-guinea pig Alexa Fluor 488; and anti-chicken Alexa Fluor 647 (all from Thermo Fisher Scientific).

To identify which receptor(s) mediate the effects of CHI3L1, we performed RNA interference-mediated knockdown of candidate receptors in iNPCs or iOPCs. Validated siRNAs targeting *CD44*, *CRTH2*, *Gal3*, *IL13Rα2*, *RAGE*, and *TMEM219* were used, alongside a control non-targeting siRNA (siCtrl; Santa Cruz Biotechnology, #sc-37007).

Briefly, 20 nM of siRNA was diluted in Opti-MEM and complexed with Lipofectamine RNAiMAX Transfection Reagent at a 3:1 ratio (reagent:siRNA/Opti-MEM, volume:volume). The mixture was then added directly to the cell culture. Twenty-four hours post-transfection, recombinant human CHI3L1 was introduced to the culture medium. Cells were maintained in the presence of CHI3L1 for a further 2 days for proliferation assays (assessed by EdU incorporation) or for 3 days for differentiation assays (assessed by expression of DCX for neurogenesis or Olig2 for oligodendrogenesis).

To investigate the downstream cell death pathways mediating the effects of CHI3L1, we utilized pharmacological inhibitors targeting key cell death modalities. Inhibitors of pan-caspase (apoptosis), necroptosis, ferroptosis, and pyroptosis were obtained from commercial sources (MedChemExpress, MCE). For all experiments, iNPCs were pre-treated with the respective inhibitors for 2 hours prior to the addition of recombinant human CHI3L1. Cells were then cultured in the continuous presence of both CHI3L1 and the inhibitor for 3 days, after which they were fixed and immunostained for Olig2 and DCX to assess oligodendrogenesis and neurogenesis, respectively. All inhibitors were carefully selected based on their established efficacy and target specificity to ensure accurate interrogation of CHI3L1’s mechanism of action in OPC generation.

### Immunoblotting and the densitometric analysis for protein levels

Protein lysates were prepared from iNPC monocultures or iNPC/iAstro cocultures using RIPA lysis buffer supplemented with protease and phosphatase inhibitors. Equal amounts of protein were resolved by SDS-PAGE on 8–15% gradient gels and transferred to polyvinylidene difluoride (PVDF) membranes that had been pre-activated with methanol. Membranes were blocked with 5% nonfat dry milk in TBST for 1 hour at 37°C and then incubated with primary antibodies overnight at 4°C. After washing with TBST, membranes were incubated with horseradish peroxidase (HRP)-conjugated secondary antibodies for 1 hour at room temperature. Protein bands were visualized using enhanced chemiluminescence (ECL Plus) reagent and imaged on an iBright Imaging System. Band intensities were quantified using ImageJ software (https://imagej.net/ij/) and normalized to the corresponding loading control.

The following primary antibodies were used: rabbit anti-CHI3L1 (1:1000; Abcam, #ab77528), mouse anti-GFAP (1:1000; Cell Signaling Technology, #3670), mouse anti-PCNA (1:1000; Invitrogen, #PC10), rabbit anti-CRTH2 (1:1000; Invitrogen, #PA5-20332), and chicken anti-Olig2 (1:1000; Aves Labs, #OLIG2-0100). For cell death and autophagy analyses, an antibody sampler kit (Bio-Rad, #42867T) was used according to the manufacturer’s instructions, targeting cleaved caspase-3, cleaved PARP (Asp214), phospho-RIP (Ser166), phospho-RIP3 (Ser227), phospho-MLKL (Ser358), cleaved gasdermin D (Asp275), LC3B, and SQSTM1/p62. Mouse anti-β-actin (1:1000; Sigma-Aldrich, #A197) or rabbit anti-vinculin (VCL; 1:1000; Invitrogen, #42H89L44) served as loading controls.

### Chemicals

Acetylcysteine (N-Acetylcysteine, MCE: HY-B0215), Azeliragon (TTP488, MCE: HY-50682), Deferoxamine Mesylate (MCE: HY-B0988), Aspartic acid (MCE: HY-42068), FPS-ZM1 (MCE: HY-19370), 1,5-Isoquinolinediol (MCE: HY-W015422), Z-VAD-FMK (MCE: HY-16658B), Necrostatin-1 (MCE: HY-15760), Ferrostatin-1 (MCE: HY-100579), Disulfiram (Tetraethylthiuram disulfide, MCE: HY-B0240) were incubated in culture as indicated.

### q-PCR

mRNA was harvested with the Qiagen RNeasy Mini Kit (Qiagen #74104). cDNA was synthesized following the iScript™ gDNA Clear cDNA Synthesis Kit (Bio-Rad #1725035). SYBR green probes (obtained from Bio-Rad) and FAM probes (obtained from iDT) targeting genes of interest were applied to evaluate gene expression under the CFX96 Touch Real-Time PCR Detection System. The gene expressions relative to loading (GAPDH/TUBB3) were normalized to the control for comparison.

### Statistical analysis

The normality of the data distribution was routinely determined by a Shapiro-Wilk normality test (p < 0.05 indicating a nonnormal distribution). For the data confirmed to be normally distributed data, we used Student t test for pairwise comparisons and one-way or two-way analysis of variance (ANOVA) followed by Tukey’s post hoc test for three groups or more, as indicated in the figure legends. For data that are not normally distributed, nonparametric alternatives, such as Mann-Whitney or Kruskal-Wallis tests, were used. All data in bar graphs and summary plots were shown as means ± SEM of at least three independent biological replicates in all figures. Significance was considered if * p< 0.05, ** p< 0.01, *** p< 0.001, and **** p< 0.0001. All statistical were performed in GraphPad Prism 9 (GraphPad Software).

### Data and materials availability

All data associated with this study are present in the paper or the Supplementary Materials. The complete raw numeric data underlying all figures, including individual data points, group summaries, and detailed statistical analyses, have been deposited in Zenodo

## Acknowledgments

This project was supported by grants from National Institutes of Health (R01AG083943) and Alzheimer’s Association (ABA-22-965518) to Dr. Yu-wen Alvin Huang.

## Author Contributions

X.Y. and Y.A.H. conceived and designed the study. X.Y. performed all experiments, analyzed the data, and prepared the figures. Y.A.H. acquired funding. X.Y. and Y.A.H. wrote the manuscript. All authors discussed the results and contributed to the review and editing of the final manuscript.

## Declarations

Y.A.H. is a founder of Acre Therapeutics LLC, a company focused on the development of antisense oligonucleotide (ASO) therapies for tauopathies, including Alzheimer’s disease. The remaining authors declare no competing interests.

## Declaration of generative AI and AI-assisted technologies in the writing process

During the preparation of this work the authors used Hemingway Editor in order to correct grammar errors and minor language issues for the first draft. After using this tool/service, the authors reviewed and edited the content as needed and take full responsibility for the content of the publication.

---

**Figure S1.**
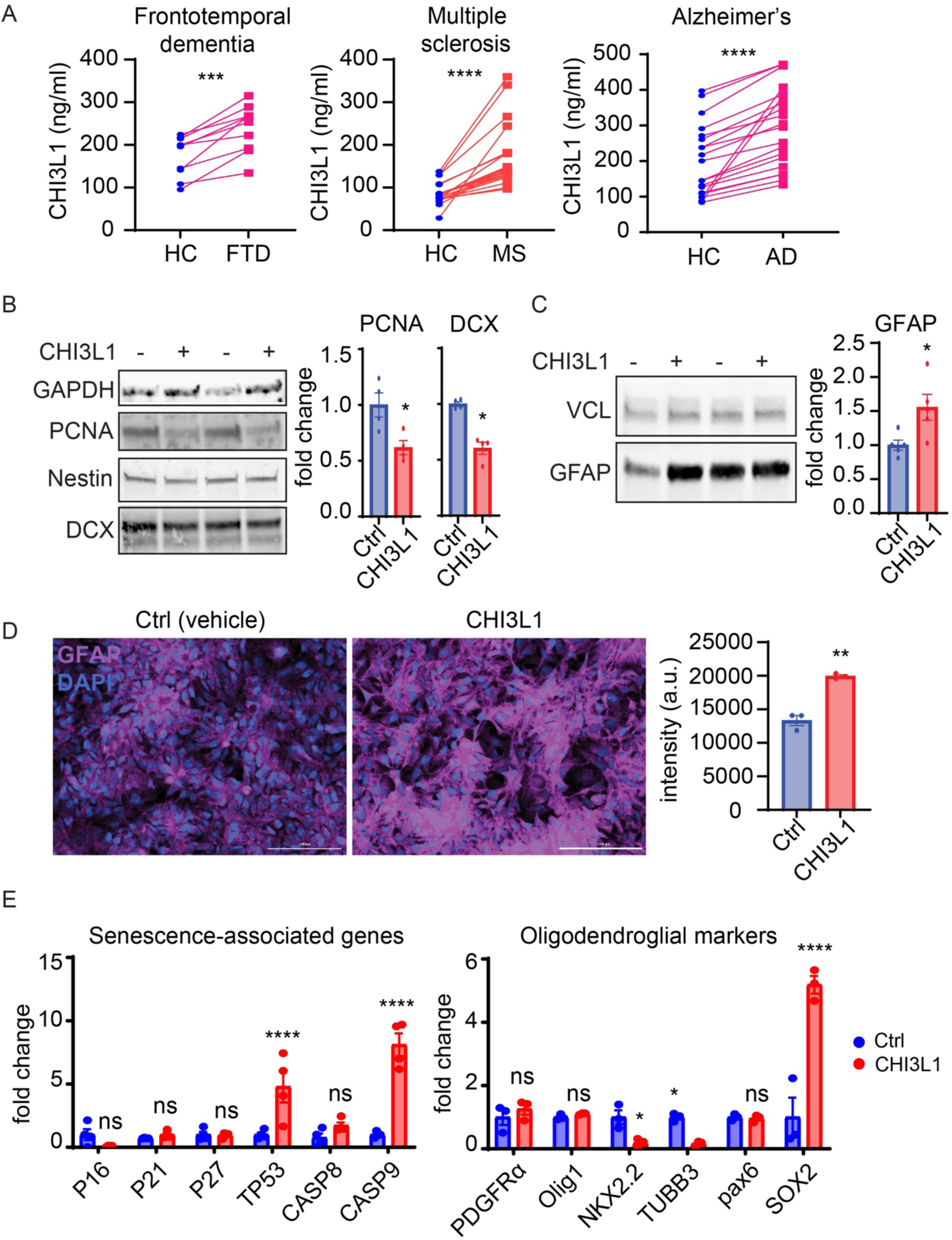
CHI3L1 impairs neural progenitor proliferation and differentiation and is elevated in neurological disease states (related to Figure 1) **(A)** Cerebrospinal fluid (CSF) concentrations of CHI3L1 in healthy controls (HC) and patients with frontotemporal dementia (FTD), multiple sclerosis (MS), or Alzheimer’s disease (AD), compiled from published studies. Paired comparisons show increased CHI3L1 levels across disease cohorts. *n* = 10, 23, and 20 pairs of independent reports for FTD, MS, and AD, respectively. **(B)** Immunoblotting analysis of proliferation and neurogenesis markers in differentiating human iPSC-derived neural progenitor cells (iNPCs) with or without CHI3L1 treatment. Representative immunoblots for PCNA and DCX are shown at left, with quantification at right. PCNA levels were normalized to GAPDH, and DCX levels were normalized to nestin. CHI3L1 reduced both proliferation and neuronal differentiation. *n* = 4 blots from 3 independent experiments. **(C)** Immunoblotting analysis of GFAP expression in differentiating iNPCs with or without CHI3L1 treatment. Representative immunoblots are shown at left, and quantification normalized to vinculin (VCL) is shown at right. CHI3L1 increased GFAP protein expression. *n* = 4 blots from 3 independent experiments. **(D)** Immunofluorescence staining of GFAP in differentiating iNPCs under control and CHI3L1-treated conditions. Representative images are shown at left, and quantification of GFAP fluorescence intensity is shown at right. CHI3L1 increased GFAP immunoreactivity. *n* = 3 independent experiments. **(E)** qPCR analysis of gene expression in differentiating iNPCs following CHI3L1 treatment. The left panel shows senescence- and cell death-associated genes, and the right panel shows genes associated with oligodendroglial and neural progenitor lineage progression. CHI3L1 increased expression of TP53 and CASP9, reduced expression of NKX2.2 and TUBB3, and increased SOX2, with no significant changes in several other markers. *n* = 3–4 independent experiments. Data are presented as mean ± SEM. Statistical significance was determined by paired *t* test in (A), unpaired *t* test in (B-D), and one-way ANOVA with post-hoc analyses with selected comparisons (Ctrl vs. CHI3L1) shown in (E). * p < 0.05, ** p < 0.01, **** p < 0.0001; ns, not significant.

**Figure S2.**
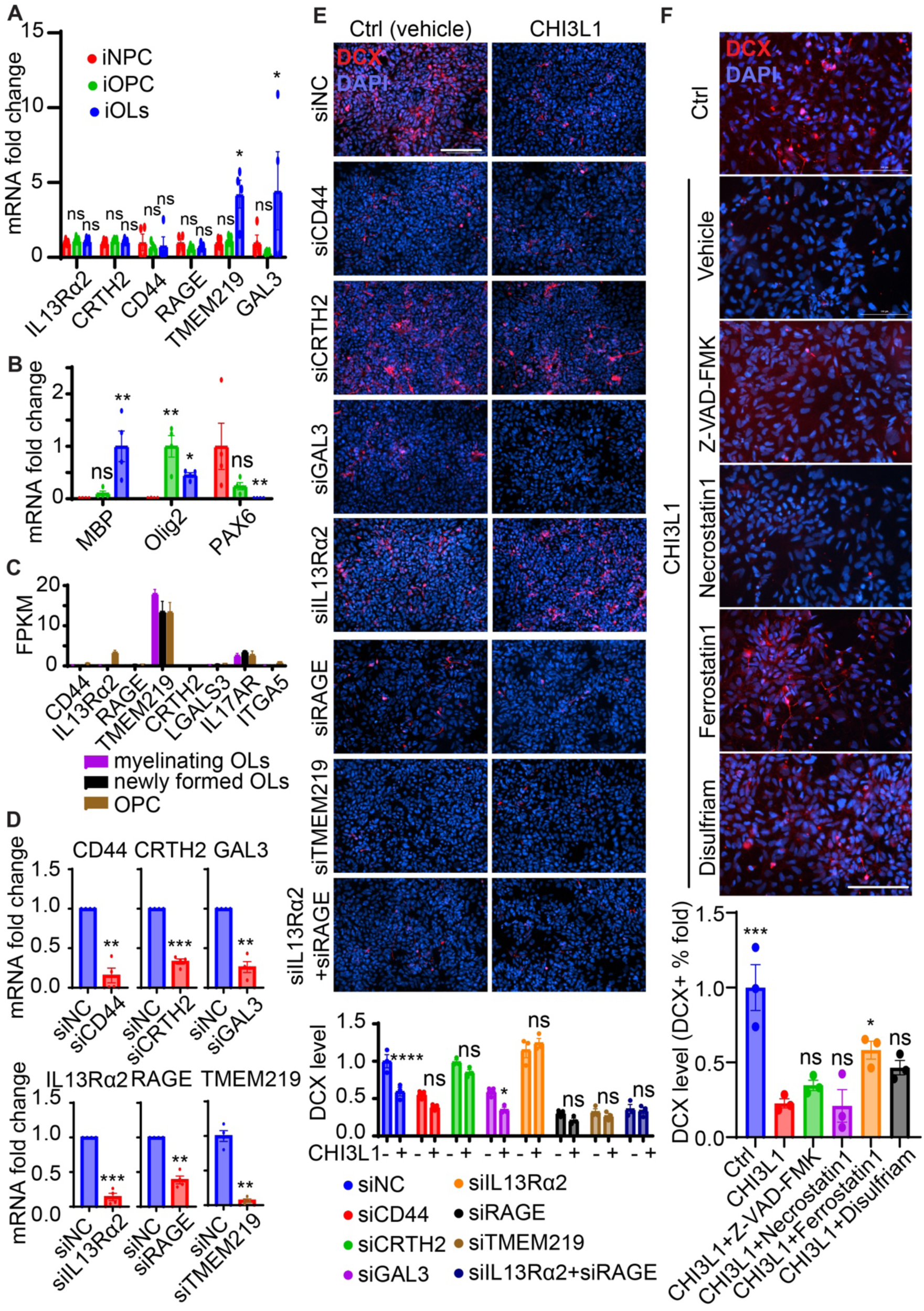
Expression and functional validation of CHI3L1 receptors in neural lineage cells and their roles in CHI3L1-mediated effects on neurogenesis (related to Figure 2) **(A)** qPCR analysis of CHI3L1 receptor expression (IL13RA2, CRTH2, CD44, RAGE, TMEM219, and LGALS3) in human iPSC-derived neural progenitor cells (iNPCs), induced oligodendrocyte precursor cells (iOPCs), and induced oligodendrocytes (iOLs). Relative mRNA levels are shown as fold change. *n* = 3 independent experiments. **(B)** qPCR analysis of lineage identity markers across differentiation stages. Expression of PAX6 (NPC marker), Olig2 (OPC/oligodendroglial lineage marker), and MBP (mature oligodendrocyte marker) confirms stage-specific differentiation of iNPCs, iOPCs, and iOLs. *n* = 3 independent experiments. **(C)** Expression profiling of CHI3L1 receptors in mouse oligodendroglial lineage populations based on an independent bulk RNA-seq dataset. Expression levels (FPKM) are shown for OPCs, newly formed oligodendrocytes, and myelinating oligodendrocytes. **(D)** Validation of siRNA-mediated knockdown efficiency for individual CHI3L1 receptors in differentiating iNPCs, assessed by qPCR. Knockdown of each target receptor significantly reduced its corresponding transcript level relative to control (siNC). *n* = 3 independent experiments. **(E)** Immunofluorescence staining of DCX (red) in differentiating iNPCs following siRNA-mediated knockdown of the indicated CHI3L1 receptors (siNC, siCD44, siCRTH2, siGAL3, siIL13RA2, siRAGE, siTMEM219, and combined siIL13RA2+siRAGE) in the presence or absence of CHI3L1. Nuclei were counterstained with DAPI (blue). Representative images are shown, with quantification of DCX levels presented below. Knockdown of CRTH2, but not IL13RA2, rescues CHI3L1-induced suppression of neurogenesis. *n* = 3 independent experiments. **(F)** Immunofluorescence staining of DCX (red) in differentiating iNPCs treated with CHI3L1 in the presence of indicated cell death inhibitors, including Z-VAD-FMK (apoptosis inhibitor), necrostatin-1 (necroptosis inhibitor), ferrostatin-1 (ferroptosis inhibitor), and disulfiram (pyroptosis inhibitor). Nuclei were counterstained with DAPI (blue). Representative images are shown, with quantification of DCX-positive cells at bottom right. Ferrostatin-1 partially restores DCX expression under CHI3L1 treatment. *n* = 3 independent experiments. Data are presented as mean ± SEM. Statistical significance was determined by unpaired t test in (D) or by one-way ANOVA with post-hoc analyses with selected comparisons shown (vs. control conditions) in (A, B, C, E and F); * p < 0.05, ** p < 0.01, *** p < 0.001, **** p < 0.0001; ns, not significant.

**Figure S3.**
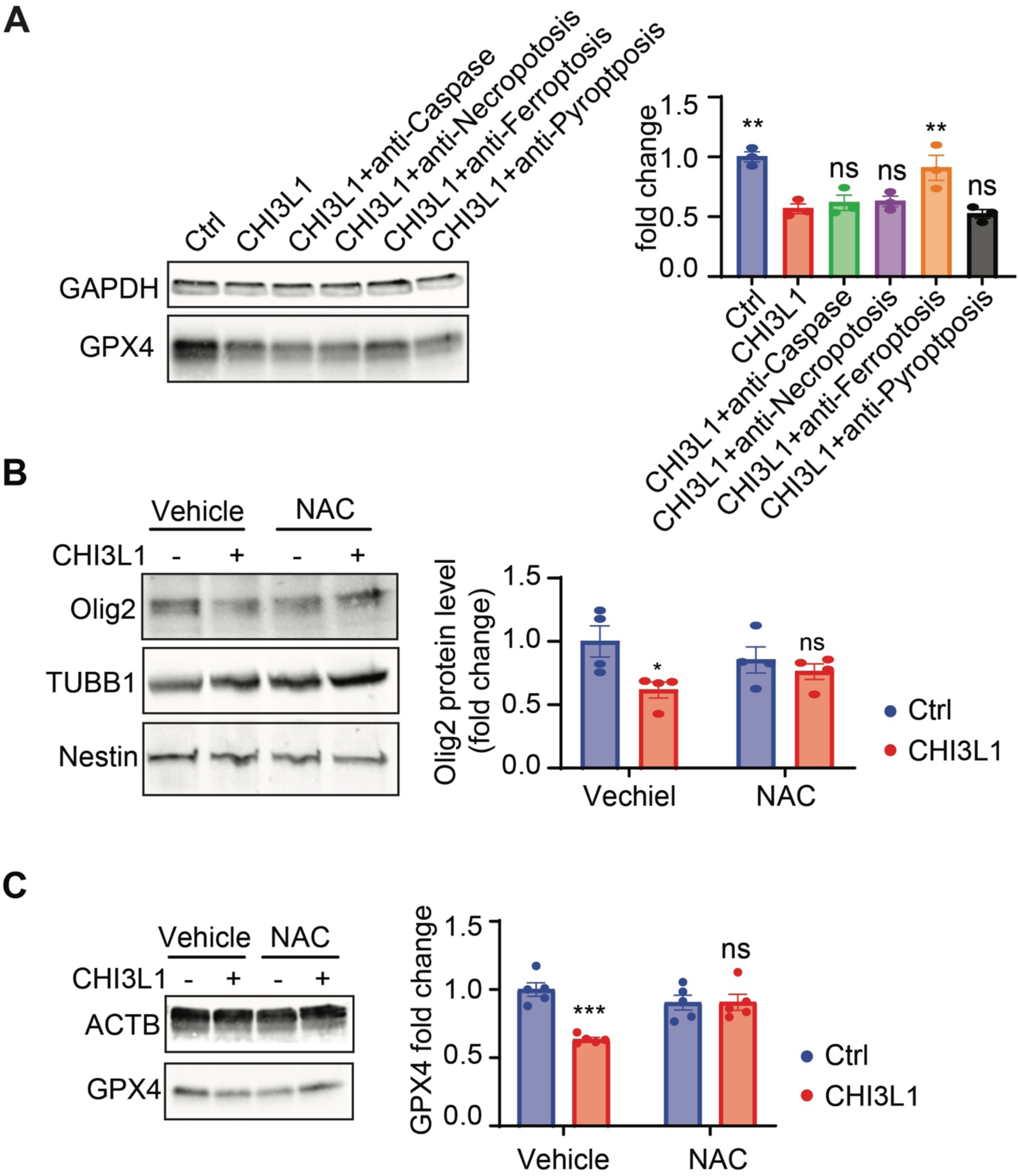
Antioxidant rescue and ferroptosis specificity in CHI3L1-mediated GPX4 suppression (related to Figure 3) **(A)** Immunoblotting analysis of GPX4 expression in differentiating human iPSC-derived neural progenitor cells (iNPCs) pre-treated with inhibitors of apoptosis (anti-caspase), necroptosis (anti-necroptosis), ferroptosis (anti-ferroptosis), or pyroptosis (anti-pyroptosis), in the presence or absence of CHI3L1. Representative immunoblots are shown at left, with quantification normalized to GAPDH shown at right. Ferroptosis inhibition selectively restores GPX4 levels following CHI3L1 treatment. *n* = 3 independent experiments. **(B)** Immunoblotting analysis of Olig2 expression in differentiating iNPCs pre-treated with the antioxidant N-acetylcysteine (NAC), with or without CHI3L1 treatment. Representative immunoblots are shown at left, with quantification normalized to the loading control shown at right. NAC attenuates CHI3L1-induced reduction of Olig2. *n* = 4 independent experiments. **(C)** Immunoblotting analysis of GPX4 expression in differentiating iNPCs pre-treated with NAC, with or without CHI3L1 treatment. Representative immunoblots are shown at left, with quantification normalized to β-actin (ACTB) shown at right. NAC restores GPX4 levels following CHI3L1 treatment. *n* = 4 independent experiments. Data are presented as mean ± SEM. Statistical significance was determined by one-way ANOVA with post-hoc analyses and the selected comparisons shown (vs. control conditions in blue bars); * p < 0.05, ** p < 0.01, *** p < 0.001, **** p < 0.0001; ns, not significant.

